# Efficient genetic code expansion tools enable *in vivo* study of lysine acetylation in non-model bacteria

**DOI:** 10.1101/2025.04.14.648644

**Authors:** Elise M. Van Fossen, Molly Stephenson, Andrew Frank, Soujanya Akella, Rowan Wooldridge, Andrew Wilson, Youngki You, Ernesto Nakayasu, Rob Egbert, Joshua Elmore

## Abstract

Recent proteomic advancements have revealed widespread Nε-lysine acetylation in pathways governing pathogenicity, metabolism, and antibiotic resistance in bacteria. The spontaneous, non-specific nature of this modification in prokaryotes obscures its biological role, necessitating prokaryotic specific *in vivo* interrogation systems. Genetic Code Expansion (GCE) offers a powerful method to investigate the roles and regulation dynamics of acetyl-lysine *in vivo* with the precise incorporation of a suite of non-canonical amino acids, including acetyl-lysine analogs. However, its use has been largely restricted to *E. coli* strains due to challenges associated with implementation and optimization of the technology in more diverse bacterial strains. Here, we present a bacterial host-agnostic, readily optimizable GCE platform designed to site-specifically incorporate non-canonical amino acids into target proteins within living bacteria. We further demonstrate the versatility of this technology by showcasing, for the first time, the successful incorporation of acetyl-lysine in a non-*E. coli* bacterium.

## INTRODUCTION

Over the past decade, it has become apparent that PTMs play crucial physiological roles in microbial organisms^1, 2^. The establishment of a comprehensive biological framework that encompasses the characteristics, regulatory mechanisms, and functions of bacterial PTMs, will unveil new opportunities for bioengineering and the treatment of infectious diseases^2^. Yet, PTMs remain one of the largest black boxes in our understanding of bacterial physiology^1–4^.

Nε-lysine acetylation is one such modification that has recently emerged as a potentially significant player in bacterial physionlogy^5–7^. Once considered rare in bacteria, mass spectrometry advances have now enabled high-sensitivity analyses of bacterial proteomes, revealing extensive lysine acetylation (up to 40% of all bacterial proteins) in more than 30 different bacterial species^8–11^. Proteomic^5, 10, 12–15^ and *in vitro* studies^5, 16–21^ have connected lysine acetylation to key biological functions, including the regulation of central metabolism^5, 6, 22^, transcription^6, 7, 23^ and translation^24^ as well as modulating pathogeneticity^25^ and antibiotic resistance^26^ (Fig. 1a). The pervasive presence of acetyl-lysine in these systems indicates that microbial bioengineering efforts and antibacterial designs that are naïve to endogenous acetylation mechanisms may exhibit diminished efficiency and efficacy upon implementation. Further, the systems that govern acetylation themselves are likely to be powerful targets for engineering enhancement and therapeutic intervention.

**Fig. 1.**
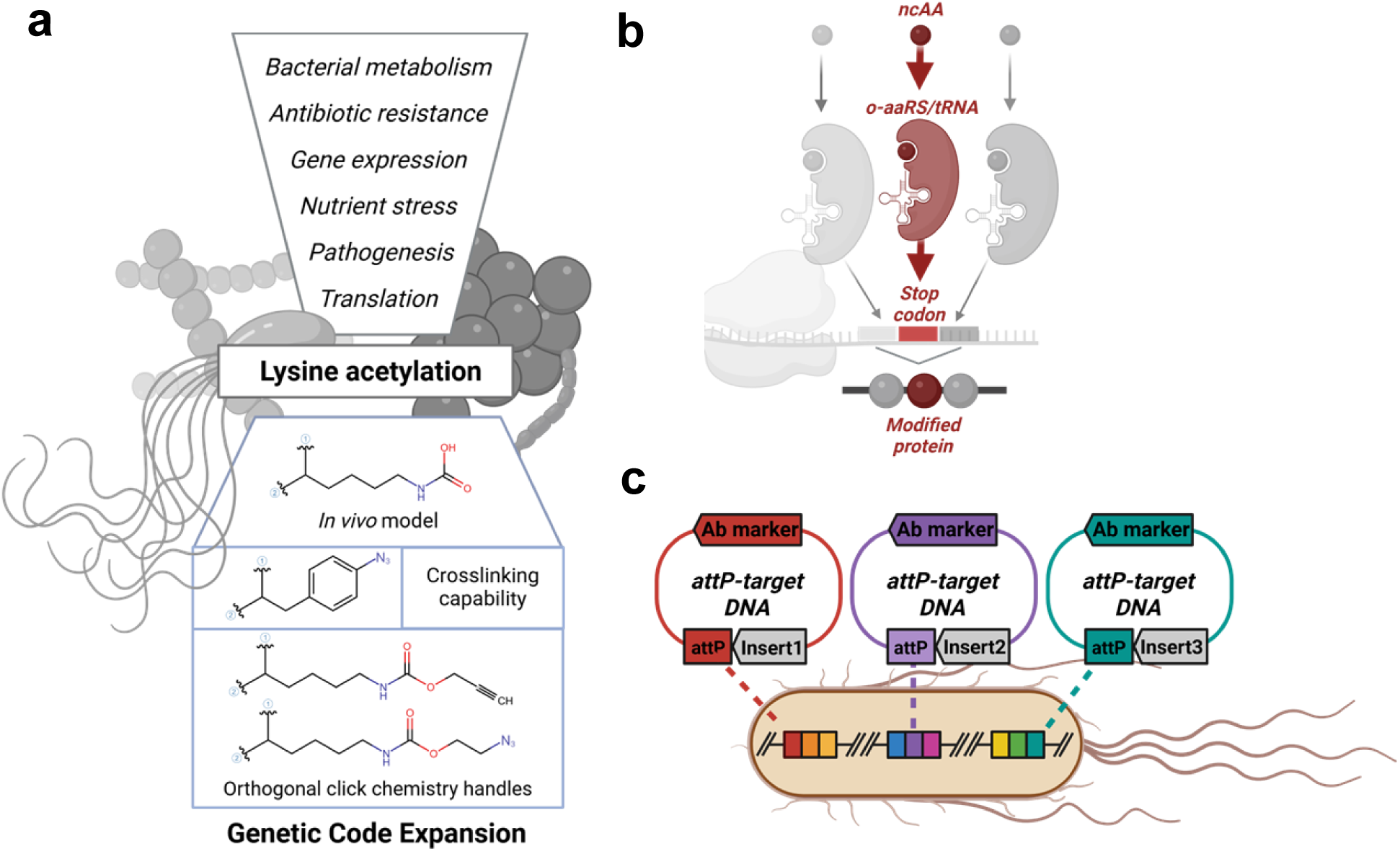
Overview of systems employed in this work. **a** Overview of lysine acetylation in diverse bacteria. Included are the chemical structures of ncAAs employed in acetyl lysine interrogation. **b** Overview of Genetic Code Expansion (GCE). Elements in red represent engineered translational components that act orthogonally to endogenous translation to produce site-specifically modified protein. **c** Overview of Serine-recombinase Assisted Genome Engineering (SAGE) where serine-recombinases encoded on non-replicating plasmids facilitate the insertion of *attP*-containing target plasmids into cognate *attB*-sequences that have been installed into the host chromosome.

Our understanding of lysine acetylation in bacterial systems remains limited due to several complicating factors. Firstly, lysine acetylation systems in bacteria differ greatly from eukaryotic systems and even vary widely among bacterial species, making it difficult to create generalized models for PTM mechaniscs^2, 7^. Secondly, lysine acetylation occurs at very low levels on numerous proteins and often in response to specific environmental conditions that can be challenging to consistently produce in model strains in typical laboratory conditions^6^. Additionally, because a significant proportion of lysine acetylation in bacteria occurs non-enzymatically and bacterial acetylases themselves are promiscuous, specialized biochemical tools are required to create acetyl-lysine mimics for analysis^7, 8^. In summary, little is known about acetyl-lysine regulatory mechanisms and interaction systems outside of *E. coli*, and what is known in *E coli* may not translate to other bacteria.

Consequently, identifying the functional roles of lysine acylation and pinpointing the mechanisms that drive it is among the main challenges in bacterial biochemistry today^7^. Doing so depends on the establishment of two key capabilities: 1) to model the presence/absence of the PTM on a target protein, and 2) to reliably characterize the interacting entities or phenotypic fluctuations associated with each state^27^. Historically, most acetylation studies have used site-directed mutagenesis to substitute the lysine residue of interest with glutamine or arginine. These substitutions mimic acetyl-lysine and non-acetylated lysine, respectively^27^.

While these studies have formed the basis of our understanding of lysine acetylation, this method has severe limitations as the substitutions replicate the electrostatic properties of the modifications but not their steric attributes. Consequently, there are numerous instances where this mutation does not accurately recapitulate the functional effects of acetyl-lysine^28, 29^.

Advancements in synthetic and chemical biology have provided new technologies with which to augment lysine interrogation toolkits. Genetic code expansion (GCE) is one such technology, wherein engineered organisms can incorporate non-canonical amino acids (ncAAs), including PTM mimics, site-specifically into proteins, allowing the precise recapitulation of lysine acetylation *in vivo* without relying on enzymatic action or site-directed mutagenesis. GCE technology has also provided ncAAs with unobtrusive crosslinking functionalities that facilitate the enrichment of and identification of PTM-driven protein-protein interactions (Fig. 1a)^27^.

As acetyl-lysine occurs in response to specific environmental conditions and is strain-specific, bacterial acetyl-lysine functional investigations should proceed in living bacterial organisms to achieve a more accurate and biologically relevant understanding^7, 23, 30^. *In vivo* application of GCE to study bacterial PTMs is nascent but has already been impactful. In *Salmonella typhimurium*, the genetically encoded incorporation of a non-hydrolysable butyryl lysine analogue into HilA (an important transcriptional regulator of *Salmonella* pathogenicity) demonstrated the consequences of specific butyrylation on infectivity^30^ and, very recently, *in vivo* GCE was employed in *E. coli* to determine how acetylation modulates transcription factor DNA binding^23^.

These two examples highlight the significant potential that *in vivo* genetic code expansion could offer for advancing our understanding of bacterial biology, however the technology still faces a critical limitation; most development and optimization of GCE has been constrained to laboratory strains of *E. coli*. As a result, when working with more diverse bacteria, the efficacy of GCE elements must be determined through trial and error and there are few, if any, universal methods for optimizing GCE in non-*E. coli* strains. This limitation disproportionately impacts systems aimed at incorporating non-canonical amino acids with more subtle modifications, such as acetyl-lysine, as the incorporation efficiency for these smaller modifications tends to be lower compared to ncAAs with bulkier side chains^31^. Consequently, optimizing these systems to perform robust science in non-model bacterial hosts remain technically challenging and laborious.

The lack of high-throughput genetic tools for the rapid optimization of expression levels of GCE machinery is a major bottleneck to onboarding complex GCE systems in non-model microbes. Common methods for transferring heterologous DNA into non-model bacteria— replicating plasmids, homologous recombination-based allelic exchange, and transposon mutagenesis—have limitations that restrict their utility. Replicating plasmids are often unstable, impose a fitness cost, and have a limited number of compatible plasmid options^32, 33^. While stable integration of heterologous DNA via transposon-based and homologous recombination-based technologies can bypass some of these problems, these methods are not suitable for high-throughput genetic engineering^34^. Transposon-based tools have unpredictable integration sites, risking overestimation of GCE efficiency due to the influence of local environment on machinery expression^35^. Homologous recombination, being low-efficiency and labor-intensive, is unsuitable for high-throughput assessments of translational machinery expression variants^36^.

However, the genetic engineering platform SAGE (Serine recombinase-assisted genome engineering) and similar phase recombinase-based systems offer the potential to overcome these limitations by combining the high-efficiency transformation of replicating plasmids, with the site specificity and stability of homologous recombination. SAGE is a robust and extensible technology that enables site-specific genome integration of multiple DNA constructs, often with efficiency on par with or superior to replicating plasmids^35^. The toolkit leverages high-efficiency serine recombinases, each transiently expressed from a non-replicating plasmid, to facilitate efficient, iterative integration of constructs or libraries into diverse bacterial genomes at unique attB sites (Fig. 1c). By utilizing non-replicating plasmids, SAGE is theoretically usable in any bacterial species and has been demonstrated in eight taxonomically distinct bacteria thus far. As such, in this study we apply SAGE to implement GCE in five non-model bacteria to demonstrate a generalizable workflow for onboarding and optimizing genetic code expansion in organisms that are unrelated to *E. coli*. Using this workflow, we dramatically enhanced the efficiency of acetyl-lysine incorporation in non-model bacteria—from nearly negligible levels to a robust system capable of encoding acetyl-lysine at two distinct sites within a biologically relevant protein.

## RESULTS

### Multiplexed SAGE enables facile application of genetic code expansion in Pseudomonas putida KT2440

Implementation of plasmid-free genetic code expansion requires the installation of an orthogonal translation system composed of, at minimum, three genetic elements into the chromosome of the host organism. These components include the <u>o</u>rthogonal <u>a</u>mino<u>a</u>cyl t<u>R</u>NA <u>s</u>ynthetase (o-aaRS), its cognate <u>o</u>rthogonal tRNA (o-tRNA) and a target protein. This target protein contains a ‘blank’ codon (typically a stop codon) at the desired site of ncAA incorporation (Fig. 1b). The orthogonal elements have been engineered to recognize and “suppress” the stop codon with the ncAA to create full length protein^37^. For our initial evaluation of SAGE-GCE, we decided to pursue the incorporation of the ncAA *para*-azido-L-phenylalanine (pAzF) with the *M. jannaschii* tyrosyl-tRNA synthetase/tRNA^Tyr^ (*Mj*TyrRS/tRNA^Tyr^) system, specifically using the *para*-cyano-L-phenylalanine aminoacyl-tRNA synthetase (*p*CNF-RS)^34^. This system is a standard for GCE implementation in bacterial systems, as it is highly efficient and polyspecific with the ability to recognize at least 18 ncAAs, including *p*AzF, *p*CNF, and other *para*-substituted phenylalanine analogues^34^. Additionally, pAzF is a valuable ncAA to encode as it contains a chemical handle compatible with crosslinking experiments, applicable for the identification of PTM-dependent protein-protein interactions^27^.

Genetically incorporating each component into individual attB sites provides the ability to independently optimize each component. For this, we utilized a variation of SAGE to incorporate multiple distinct ‘target plasmids’ into the chromosome without the need to excise the antibiotic selection marker between each incorporation as is required with the original SAGE implementation (Fig. 1c). For this, each component is linked with one of three antibiotic selection and serine recombinase combinations, which enables incorporation of each element into the chromosome in any order. For SAGE-GCE, we designed target plasmids containing each of the three required components with unique antibiotic resistance markers. These plasmids also included distinct *attP* attachment sites for integration into three distinct *attB* sites in the bacterial chromosome by a set of three serine recombinases. Specifically, the *Mj*TyrRS element was designed for insertion at the TG1 *attB* site, the tRNA^Tyr^ element at the Bxb1 *attB* site, and the protein of interest, sfGFP in this case, at the R4 *attB* site (Fig. 2a).

**Fig. 2.**
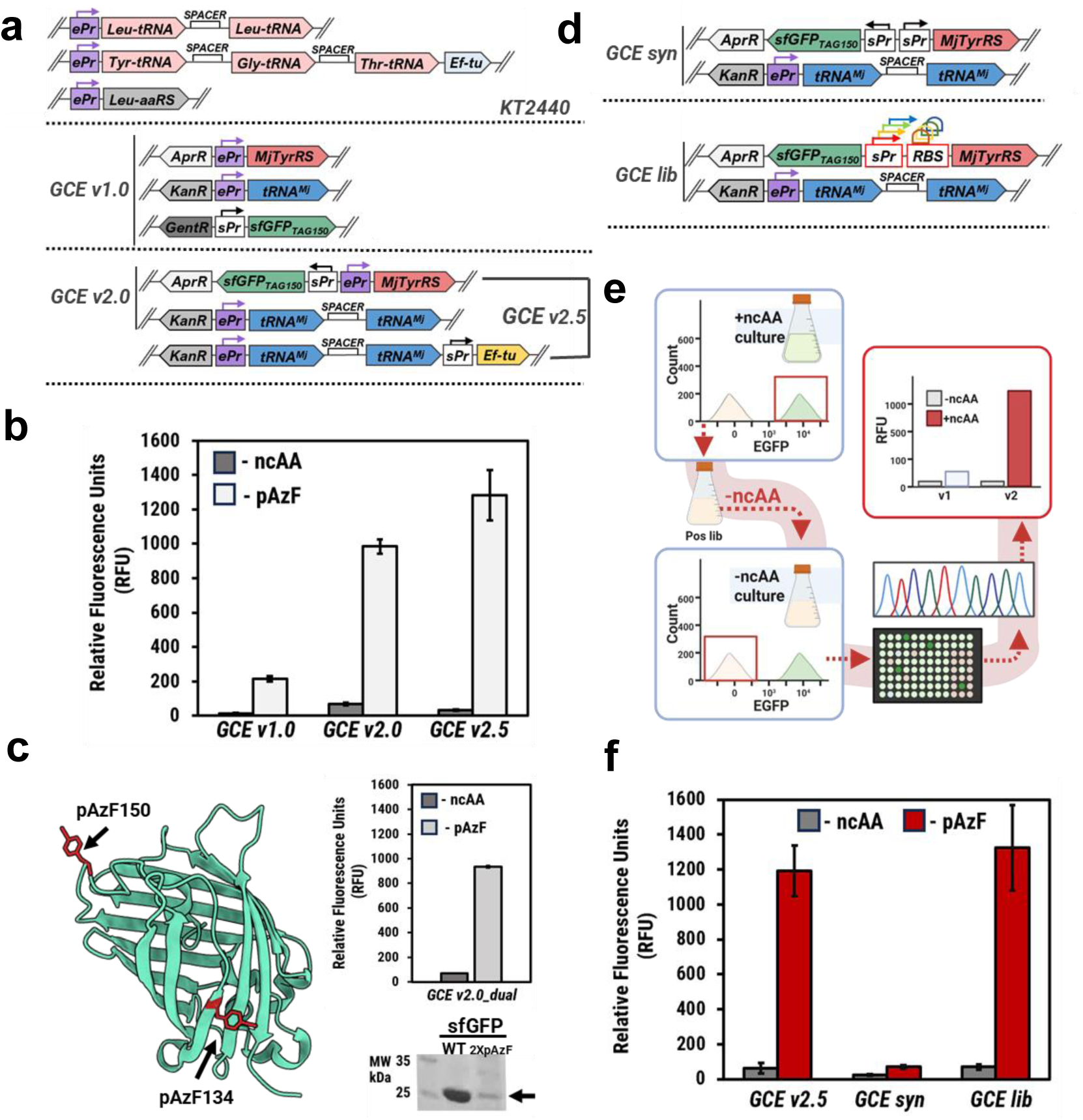
GCE optimization strategies in P. putida. **a** Overview of endogenous and initial GCE translational systems utilized in this work. **b** GCE efficiency assays for KT2440 GCE-SAGE systems from four biologically independent cultures. The GCE system used is indicated below each column set on the *x*-axis. The ncAA condition for each system is disclosed in a legend in the upper left of the figure. **c** Overview of dual incorporation system. Structure of sfGFP is indicated on the left with each site indicated in red. GCE incorporation efficiency for the dual incorporation system is shown on the right from large scale expression cultures of KT2440. SDS-PAGE of 2X pAzF shown side-by-side with WT GFP, protein band of interest is indicated by an arrow. **d** Overview of GCE synthetic translational systems and GCE library scheme. **e** Scheme of FACS-based library selection for promoter optimization. After transformation into KT2440, the promoter library is cultured in the presence of ncAA to elicit a fluorescent signal. Fluorescent events are sorted via FACS and cultured in the absence of ncAA. Low fluorescent events are then plate sorted for sequencing and final validation. **f** GCE efficiency assays for KT2440 GCE synthetic translational system and optimized GCE library system from four biologically independent cultures. The ncAA condition for each system is disclosed in a legend at the top of the figure.

Our initial test platform for the SAGE-based GCE system was the bacterium *Pseudomonas putida* KT2440^38, 39^. *P. putida* has become increasingly popular for industrial and environmental applications^40^ due to its robust redox metabolism, high tolerance to diverse physicochemical stresses, rapid growth, versatile metabolism, and nonpathogenic nature. Recently, successful replicating plasmid-based GCE was reported in *P. putida* for the first time^41^ encouraging our efforts to use this system as a testbed for our SAGE-based GCE system. For this, we used a SAGE-compatible strain of *P. putida* KT2440 (AG5577) for the basis of all further GCE development and application in *P. putida*. This strain contains a collection of 9 distinct heterologous *attB* sequences that are each recognized by a distinct serine recombinase from the SAGE toolkit^35^.

Multiple studies focused on implementing GCE in non-*E. coli* strains have reported that it is critical to adjust the regulatory elements (promoters, ribosome binding sites, etc.) for each new organism^42–44^. To maintain GCE components at physiologically relevant expression levels in *P. putida*, we followed a previously reported strategy^41, 44^ wherein the most abundant codon in the *P. putida* genome was identified (CUG, encoding leucine (Leu)) along with the associated aaRS and tRNA. Native promoters and terminators flanking tRNA^Leu^ and leucyl-RS (LeuRS) were then used to control transcription of the *Mj* tRNA^Tyr^ and *Mj*TyrRS, respectively to create GCE v1.0 (Fig. 2a).

Once all SAGE-GCE components were successfully ported into *P. putida* (see Methods), ncAA incorporation efficiency was evaluated using a stop codon readthrough assay with super-folder green fluorescent protein (sfGFP) as a reporter. In this assay, sfGFP contains an amber stop codon at position N150 (sfGFP_TAG150_). In the absence of ncAA incorporation, expression of this reporter will lead to the synthesis of a truncated, non-fluorescent sfGFP protein. If the ncAA is incorporated into the protein at the amber codon, full length fluorescent sfGFP protein will be synthesized and the cellular fluorescence can be directly correlated with production of ncAA-containing sfGFP protein. Consistent with what was previously reported in *P. putida*, we observed that the pCNF-RS system enabled efficient incorporation of pAzF as indicated by moderate expression of full-length sfGFP only in the presence of the ncAA (Fig. 2b). Further top-down protein mass spectrometry (MS) analysis of the full-length purified protein confirmed the incorporation of pAzF (Fig. S1).

Although this initial GCE system exhibited successful incorporation of pAzF, we noted that the efficiency could much improved. As a high concentration of o-tRNA is required for optimal incorporation efficiency in other GCE systems^45^, we hypothesized that increasing the copies of o-tRNA in our system would enhance ncAA suppression. To test this hypothesis, we designed a tRNA cassette to mimic an endogenous *P. putida* tRNA operon, which encodes two identical tRNA^Leu^ in a single pre-tRNA transcript (Fig. 2a, GCE v2.0). This cassette utilizes the endogenous tRNA^Leu^ promoter, the spacer sequence between the tRNAs, and the native downstream terminators, but replaces each tRNA^Leu^ with a *Mj*tRNA^Tyr^.

We also observed a *P. putida* tRNA operon containing three tRNAs positioned upstream of a gene encoding an elongation factor protein (EF-Tu), an enzyme responsible for facilitating the transfer of aminoacyl-tRNA to the ribosome. As native EF-Tus can exhibit decreased efficiencies when operating in conjunction with ncAAs and orthogonal elements, we drew inspiration from the endogenous *P. putida* operon and added an engineered EF-Tu—designed specifically for the incorporation of pAzF^46^—downstream of the dual *Mj*tRNA^Tyr^ operon described above, creating GCE v2.5. Lastly, we streamlined GCE v1.0 by combining the sfGFP_TAG150_ reporter and *Mj*TyrRS on a single construct resulting in a dual plasmid system (Fig. 2a). We evaluated the incorporation efficiencies of the two new GCE systems (GCE v2.0 and GCE v2.5) and found that by adding a second copy of *Mj*tRNA^Tyr^ (GCE v2.0), we achieved a five-fold improvement compared to the original single tRNA system. The addition of the engineered EF-Tu in GCE v2.5 led to a modest further 1.5-fold improvement (Fig. 2b).

We further tested the efficiency of GCE v2.0 by designing a sfGFP construct with two sites for ncAA incorporation (sfGFP_2XpAzF_, sites D134TAG and N150TAG) to direct the dual incorporation of pAzF into sfGFP (Fig. 2c). We observed reasonable efficiency for dual incorporation with GCE v2.0, highlighting the feasibility of the system for incorporating multiple ncAAs (Fig. 2c). Top-down protein MS analysis confirmed the dual incorporation of pAzF, although the protein experienced some slight degradation during purification which was accounted for in our mass calculation (Fig. S2).

### SAGE enables high-throughput library-based GCE optimization directly in the hosting organism

Multi-component systems often require the coordinated optimization of each component for high performance^47^. A common strategy is to improve transcription and translational rates by using engineered promoters for enhanced RNA polymerase recruitment or by altering the 5′ UTR to include efficient ribosomal binding sites (RBSs). While the LeuRS promoter enabled robust performance in GCE v2.0, the absence of a clear RNA polymerase binding motif posed a challenge for engineering increased transcription. The synthetic *tac* promoter is broadly used in bacteria, provides strong expression, and is easily tunable due to its well characterized core promoter elements^48^. Therefore, we evaluated the *tac* promoter as a replacement for the LeuRS promoter to support *Mj*TyrRS expression for efficient ncAA incorporation in *P. putida* (GCE syn system, Figs. 2d and 2f). Unexpectedly, sfGFP_TAG150_ expression significantly decreased when *Mj*TyrRS transcription was driven by the tac promoter compared to the endogenous LeuRS promoter (Fig. 2f). However, this outcome aligns with observations by others that tuning the expression of GCE machinery that was originally developed for use in *E. coli* is crucial for robust GCE application in non-*E. coli* strains^42^.

Rather than laboriously evaluating many promoter and ribosomal binding site variants individually to identify alternative promoters that enable improved performance, we leveraged the high transformation efficiency afforded by SAGE and the amenability of the sfGFP reporter assay to evaluate a large collection of *Mj*TyrRS expression variants in a pooled assay. We previously demonstrated the efficacy of SAGE for high-throughput sequencing-based methods to assess large promoter libraries in pooled assays^35^. Adopting a similar approach, we aimed to optimize *Mj*TyrRS production by identifying regulatory elements that ensure sufficient *Mj*TyrRS levels for robust ncAA incorporation without causing non-specific amino acid incorporation.

We constructed a pooled *Mj*TyrRS expression library using 449 synthetic and natural promoter sequences (Table S1). To achieve a diverse range of expression levels, we employed a cloning strategy to randomly incorporate one of 106 ribosomal binding site elements downstream of the promoter sequences. These RBS elements were designed by a RBS calculator^49^ to span approximately a 441-fold range of translational efficiencies (Table S2). Additionally, each regulatory sequence included the riboJ insulator sequence^50^ between the promoter and ribosomal binding site to stabilize gene expression. The resulting ∼209,000 member *Mj*TyrRS expression plasmid library was integrated into *Pseudomonas putida* using SAGE, resulting in a pooled collection of ∼10,000,000 transformants, each containing a distinct promoter-RBS combination driving *Mj*TyrRS expression. Each transformant also contained the reporter construct and the dual o-tRNA cassette from GCE v2.0 enabling GCE (Fig. 2d).

As the read-out for GCE efficiency in our system is sfGFP, fluorescence activated cell sorting (FACS) was performed on the strain library to identify efficient and orthogonal library members. This was done by first culturing the strain library in the presence of the ncAA to allow for ncAA-dependent sfGFP production. Members that exhibit efficient ncAA incorporation and thus generate a fluorescent output are enriched via FACS (positive sort). Enriched members are then cultured in the absence of ncAA to identify members with low background incorporation of native amino acids, ensuring that orthogonality is maintained with the new regulatory elements (Fig. 2e).

For the initial positive selection, we cultivated the library in media containing pAzF and sorted the top 1.5% most fluorescent library members into a pooled library of strains. This pool of fluorescent cells was sub-cultured into media lacking pAzF and after overnight incubation, was sorted again via FACS. This time, members that exhibited no fluorescent signal above background were plate sorted individually into microtiter plates, cultivated overnight, sequenced and further validated in scaled up GCE efficiency assays (Fig. S3, Table S3).

The top performing strain displayed ncAA incorporation efficiency and orthogonality that was on par with GCE v2.5 (Fig. 2f) despite lacking an engineered EF-Tu. Surprisingly, a Lactobacillus_39770 promoter with relatively low transcriptional activity in other *Pseudomonas* sp. was identified in the top performing strain^35^. Of note, this promoter was found to have between ∼24 to 92-fold lower transcriptional activity than the *tac* promoter in our prior work^35^, suggesting that poor performance with the *tac* promoter may be a consequence of excessive *Mj*TyrRS expression. The Lactobacillus_39770 promoter was coupled with a ribosomal binding site element with a relatively high predicted RBS translation initiation rate of 67067.6, on par with the RBS from our tac system. Thus, this library selection method, in one round of positive and negative sorting, produced a promoter/RBS pair with ncAA incorporation efficiency equivalent to the endogenous promoters, permitting the use of non-endogenous promoters to drive GCE technology.

### SAGE facilitates the genetic code expansion of phylogenetically distant microbes

After successful implementation of GCE in *P. putida* KT2440, we evaluated the ability to easily transfer our optimized GCE systems into related Pseudomonas sp. (Fig. 3a). These bacteria include the plant growth promoting rhizobacterium *P. fluorescens* strain SBW25^51^ and two pseudomonads that were isolated from the endosphere of *Sorghum bicolor* under drought conditions^35, 52^. Each of the two, *P. frederiksbergensis* (TBS10)^35^ and *P. facilor* (TBS28)^52^, have been previously engineered to be compatible with SAGE via the incorporation of a poly-attB landing site^35, 52^.

**Fig. 3.**
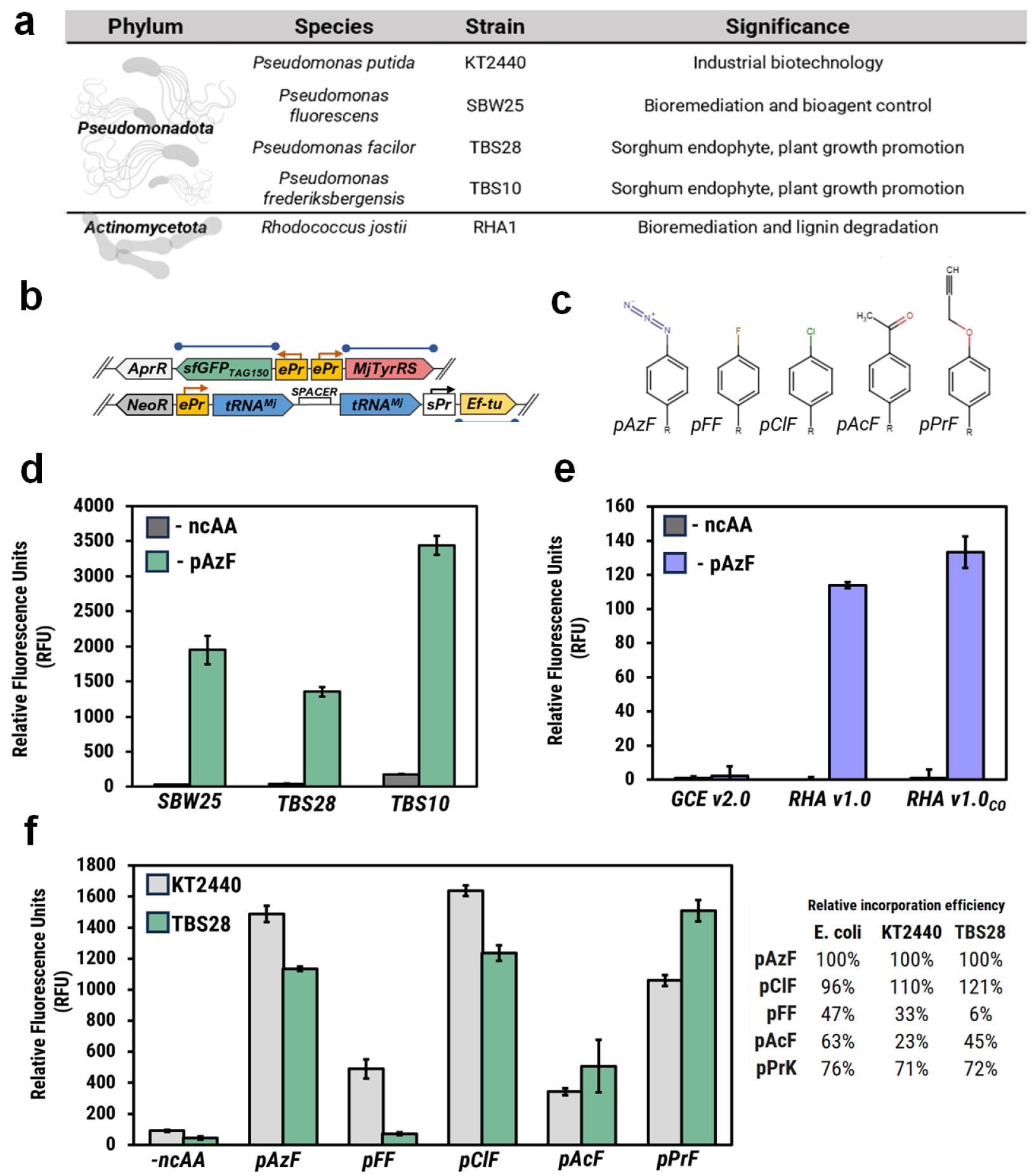
GCE-SAGE technology in non-model bacteria. **a** Overview of bacterial species used in this study. **b** Overview of RHA1 GCE translational systems. Elements that were codon optimized are indicated by a blue line. **c** Structures of ncAAs screened in KT2440 and TBS28 strains. **d** GCE efficiency assays for GCE v2.0 in three pseudomonads. The pseudomonad used is indicated below each column set on the *x*-axis. The ncAA condition for each system is disclosed in a legend in the upper left of the figure. **e** GCE efficiency assays for GCE v2.0 vs GCE RHA in RHA1. The GCE system used is indicated below each column set on the *x*-axis. The ncAA condition for each system is disclosed in a legend in the upper left of the figure. **f** ncAA incorporation efficiency assay for KT2440 and TBS28 with GCE v2.0. The ncAA system used is indicated below each column set on the *x*-axis. The strain (either KT2440 or TBS28) disclosed in a legend in the upper left of the figure. The incorporation efficiency of each ncAA relative to pAzF are shown for E. coli, KT2440 and TBS28.

We directly transferred the GCE v2.0 machinery developed in *P. putida* into the chromosomes of the three Pseudomonads and observed pAzF incorporation on par with what was observed in *P. putida* demonstrating that SAGE-based GCE systems can be shared across genetically similar bacteria (Figs. 3d and 3f). Interestingly, even though the GCE efficiency was similar across the pseudomonads, the fluorescent output did vary across a ∼2-fold range. It remains to be seen whether these differences are a consequence of gene expression differences, or of another physiological trait such as ncAA transport.

As mentioned above, the *p*CNF-RS used here is polyspecific, meaning it has the capacity to charge the o-tRNA with several distinctive *para*-substituted phenylalanine analogs. However, each organism has its own distinct set of metabolite transporters, sensors, and other physiological regulators that control expression of the proteins that enable ncAA uptake and it remains unclear how much of a role this plays in GCE efficiency. To examine this, we utilized the permissivity of the pCNF-RS employed in our GCE system^53^ to conduct a small ncAA screen, testing four additional *para*-substituted phenylalanine derivatives in both *P. putida* and the environmental isolate *P. facilor* (Fig. 3c). We then compared their incorporation efficiencies relative to pAzF to reported values from *E. coli*^53^ (Fig. 3f).

While overall the relative incorporation efficiencies among the three organisms were similar, we observed a few notable differences. Specifically, while both pseudomonads exhibited lower incorporation of 4-fluoro-L-phenylalanine relative to *E coli*, incorporation of this ncAA into sfGFP was almost non-existent in *P. facilor* (Fig. 3f). These differences are unlikely to be due to the activity of the pCNF-RS, as the other ncAAs showed very similar efficiencies. It is more likely that differences in amino acid transport between the organisms are responsible, again underscoring the importance of considering ncAA import mechanisms when integrating new bacterial strains.

A benefit of utilizing SAGE-based tools is the ability to easily transfer materials developed in SAGE-compatible hosts with phylogenetically distant SAGE-compatible bacteria. To demonstrate this benefit, we tested our GCE 2.0 system developed in *Pseudomonas putida* in the actinomycete *Rhodococcus jostii* strain RHA1^54^, a representative of a genus with diverse metabolic capabilities ranging from catabolism of cholesterol^55^ and petroleum-derived hydrocarbon^56^ to conversion of lignin-derived feedstocks into useful chemicals^57^. GCE v2.0 was ported into RHA1, yet ncAA-sensitive fluorescence was not observed above background levels (Fig. 3e) suggesting the Pseudomonad-derived machinery was incompatible with RHA1.

To optimize expression, we identified the most abundant codon (Ala) in the RHA1 genome and utilized the endogenous aaRS and tRNA promoters associated with Ala to drive the GCE elements (Fig. 3b). Considering the high GC content of Rhodococcus genomes and several strong codon biases, we also created a codon-optimized version of the protein elements to assess the impact of codon optimization on GCE efficiency. The RHA1-tailored system showed a dramatic improvement compared to GCE v2.0, with more subtle differences observed between codon-optimized (RHA v1.0_CO_) and non-codon-optimized (RHA v1.0) GCE-RHA1 systems (Fig. 3b). However, the overall fluorescent output of the RHA1 system is comparable to GCE v1.0 in *P. putida*, suggesting potential for further improvement.

### Site specific incorporation of acetyl-L-lysine and other lysine analogs

We were encouraged by our initial success in the implementation of the *Mj*TyrRS-pCNF system and sought to use our optimized platform to broaden the existing lysine PTM interrogation toolkit in non-*E. coli* strains. As the *Mj*TyrRS system primarily incorporates aromatic ncAAs, we chose to onboard the Pyl system, which enables site-specific incorporation of lysine analogues due to the structural similarity between pyrrolysine—its natural substrate—and lysine^58^. Derived from *Methanosarcina* species, the Pyl system is composed of a pyrrolysyl-aminoacyl-tRNA synthetase (PylRS) and its cognate tRNA (tRNA^Pyl^) and has enabled the incorporation of a suite of ncAAs developed to interrogate lysine PTMs^59^. From these, we chose to encode acetyl-lysine (AcK), azido-lysine (AzK) and propargyloxycarbonyl-lysine (PlK) into *Pseudomonas putida* (Fig. 4b)^59^. AcK incorporation enables direct interrogation of the influence of lysine acetylation on protein function both *in vivo* and *in vitro*^59^ and AzK and PlK contain distinct bioorthogonal click chemistry handles^60^ that can be used to probe AcK-dependent protein interactions, facilitate fluorescent labeling and in some cases, to site-specifically model ubiquitin^37^.

**Fig. 4.**
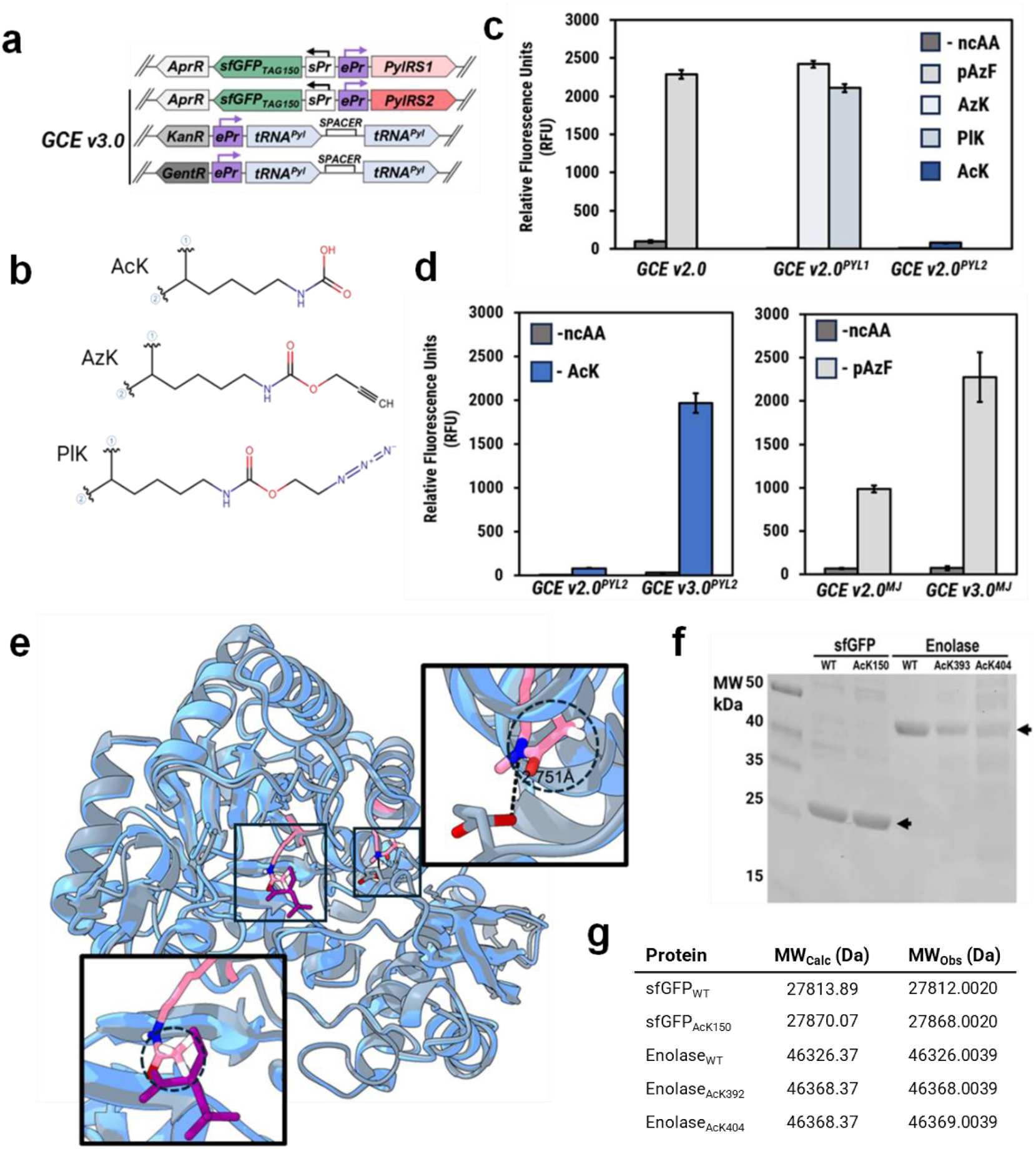
SAGE-based optimization permits facile acetyl-lysine incorporation. **a** Overview of GCE systems with Pyl components. **b** Structures of ncAAs incorporated with Pyl components. **c** GCE efficiency assays with Pyl systems in KT2440. The GCE system used is indicated below each column set on the *x*-axis. The ncAA condition for each system is disclosed in a legend on the right of the figure. **d** GCE efficiency assays comparing GCE v2.0 with GCE v3.0. Pyl system and *Mj* systems in KT2440 are compared in left panels and right panels respectively. The GCE system used is indicated below each column set on the *x*-axis. The ncAA condition for each system is disclosed in a legend in the top left of the figure. **e** Structure of enolase with acetyl lysines and bound substrate. Alpha fold generated structure of KT2440 enolase with acetylations modeled relevant lysine residues (blue ribbon structure) overlaid on PDB structure of enolase from E. coli (gray ribbon structure) with bound substrate 2PGA (purple molecule). Acetyl grounds are circled. **f** SDS-PAGE of acetylated proteins, purified from KT2440. Arrows indicate the MW for each set of purified proteins. **g** Observed intact masses of purified acetylated proteins with their respective calculated masses.

We integrated the Pyl system into *P. putida* using two engineered PylRS encased in the same genetic framework employed for GCE v2.0. The first, *Mb*PylRS-AF (PylRS1), contains active site mutations that permit the incorporation of bulky lysine analogues^61^, suitable for AzK and PlK. The second, chPylRS-IPYE (PylRS2), is a chimeric synthetase known for its high efficiency in incorporating AcK in proteins in *E. coli*^62^. These PylRSs were paired with an evolved tRNA^Pyl^, which has been shown to enhance AcK incorporation into proteins^31^ (Fig. 4a).

With PylRS1 in GCEv2.0, we observed a high incorporation efficiency of AzK and PlK with no additional alterations to the system (Fig. 4c). However, it was surprising to observe that, despite robust reported performance in *E. coli*, AcK incorporation efficiency was very low in *P. putida* (Fig. 4c). This finding, coupled with the absence of reported instances of acetyl-lysine incorporation via GCE in organisms other than *E. coli*, suggests that this ncAA poses greater challenges for incorporation than typical ncAAs, thereby restricting its application to *E. coli* systems to date.

We had previously observed that increasing *Mj*tRNA^Tyr^ copy number enhanced pAzF incorporation efficiency and posited that the tRNA^Pyl^ copy number may be a limiting factor here as well. To test this, we incorporated an additional dual tRNA operon into *P. putida* at a third *attB* site (GCE v3.0, Fig. 4a) bringing the total number of tRNA^Pyl^ copies to four. We repeated this operation for the *Mj*TyrRS-pCNF system then evaluated the ncAA incorporation for both.

Doubling the tRNA copy number improved ncAA with both systems, but to strikingly different extents. Given the already robust incorporation of pAzF with the *Mj*TyrRS system, it was unsurprising to see a substantial but relatively modest 2-fold increase in sfGFP production with the addition of the second tRNA cassette (Fig. 4d, right panel). However, the incorporation efficiency of AcK by the chPylRS(IPYE) system was improved by 25-fold (Fig. 4d, left panel), suggesting tRNA copy number was a limiting factor. This one improvement increased the efficiency of AcK incorporation to be on par with our best existing system and signified that tRNA expression is a critical lever for optimization in each host.

### Production of site-specifically acetylated enolase in Pseudomonas putida with genetic code expansion

While we were encouraged at the substantial improvement in GCE efficiency driven by additional copies of tRNA, we wanted to vet our system by producing an essential protein with acetyl-lysine encoded at biologically relevant acetylation sites. Previously, Ernesto et al performed a proteomic analysis which identified the acetylation of highly conserved lysines in central metabolic enzymes from diverse bacteria, including *Pseudomonas putida*^5^. One essential protein highlighted in the study was the enzyme enolase, which plays a crucial role in glycolysis and gluconeogenesis by catalyzing the reversible conversion of 2-phosphoglycerate to phosphoenolpyruvate, a key step in central carbon metabolism^5^. Enolase also performs additional moonlighting functions that contribute to bacterial stress responses and pathogenicity^63^. As such, we selected the enolase enzyme as a model protein to test the efficiency of AcK incorporation at multiple highly conserved lysine residues

With respect to site selection, the study identified two conserved lysines of interest in enolase; K392 which is buried within the active site and is involved in phosphoenolpyruvate catalysis and K404 which is located towards the periphery of the protein (Fig. 4e). Under the tested conditions, K404 was found to be acetylated in *Pseudomonas putida,* however, K392 which upon acetylation can ablate enolase activity, was observed to be acetylated in other bacteria but not in *Pseudomonas putida*.

*P. putida* encodes multiple lysine deacetylases, including those from the CobB and Metal-dependent lysine deacetylase families (NCBI:txid160488). Such deacetylases have a broad substrate range, but they are unable to deacetylate all acetylated lysine residues. We hypothesized that the lack of detectable acetylation at K392 may be due to efficient enzymatic deacetylation of one residue and not the other. By producing the acetylated protein *in vivo* in *Pseudomonas putida*, we could observe whether the modification is maintained at each residue over the course of a typical culturing experiment. If the acetylation remains stable, then that suggests the protein was simply not acetylated under the tested conditions.

For site-specific acetylated enolase production, we replaced the gene for sfGFP_TAG150_ in GCE v3.0 with a series of enolase constructs (enolase_WT_, enolase_TAG392_ and enolase_TAG404_) in *P. putida*. Cultures for strains harboring each of these constructs were grown to stationary phase, at which point protein was extracted and affinity-purified to assess expression and acetylation of recombinant enolase. The three variants expressed successfully based on SDS-PAGE analysis (Fig. 4F) and the intact molecular masses of all enolase variants were determined via mass spectrometric analysis. This analysis revealed masses corresponding to acetyl-lysine incorporation for enolase_TAG392_ and enolase_TAG404_ (Figs. 4G and Figs. S4, S5). While the molecular weights for other contaminating proteins were detected in the analysis, the mass associated with enolase_WT_, was not observed in the acetylated enolase samples (Fig. S5). While not exhaustive, this data refutes the hypothesis that intrinsic lysine deacetylase activity during the cultivation was responsible for the lack of acetylation at K392

## DISCUSSION

Genetic code expansion has revolutionized the analysis of the bacterial acetylome by enabling the precise incorporation of PTMs along with a suite of PTM enrichment and visualization tools, enabling real-time monitoring of acetylation dynamics under physiological conditions. However, inconsistent genetic tractability and unpredictable translational regulatory elements across microbes have hindered the implementation and optimization of this technology in diverse bacteria, particularly those of interest for human health and bioremediation. This is most evident with respect to the ncAA acetyl-lysine itself, as the subtlety of the modification makes it a more challenging incorporation target.

We address this challenge by integrating GCE with serine recombinase-assisted genome engineering (SAGE), a method renowned for its high efficiency across a wide range of bacterial strains. With GCE-SAGE, each GCE component is site-specifically integrated into the bacterial genome at a high enough efficiency for library optimization and with enough landing pads (attB sites) to integrate up to ten constructs genetically. This strategy allowed for the stable integration of GCE components into the genome of *P. putida* and improved efficiency by easily allowing for increased o-tRNA copy number. SAGE also enabled the development of a high-throughput promoter and RBS library selection method using FACS. This approach identified highly efficient promoter/RBS pairs, facilitating the fine-tuning of gene expression in non-model organisms. This technique is crucial for matching GCE output to endogenous production levels, especially when only synthetic promoters or unknown promoter strengths are available.

GCE-SAGE was then successfully transferred to multiple Proteobacteria (*Pseudomonas fluorescens* SBW25, *Pseudomonas frederiksbergensis* TBS10, and *Pseudomonas facilor* TBS28) and the Actinobacterium *Rhodococcus jostii* RHA1. Screening four para-substituted phenylalanine derivatives revealed organism-specific differences in incorporation efficiency, likely due to variations in amino acid transport rather than pCNF-RS activity. Studying and expressing amino acid transporters will be a critical component to enhance ncAA uptake as SAGE enables the application of GCE in new model systems.

We successfully incorporated lysine analogs (AzK, PrK, and AcK) using evolved pyrrolysyl-tRNA synthetases (PylRS). The variable efficiency of incorporation, particularly for AcK, highlighted the need for additional tRNA copies to achieve biologically relevant levels, confirming the critical role of tRNA abundance in GCE performance. It was encouraging that the efficiency of the system could be enhanced from near zero levels by simply adding additional copies of tRNA. As this has been observed in mammalian systems as well, it is likely that this optimization approach is near universal, providing a simple path towards GCE optimization.

The stable incorporation of acetyl-lysine at two distinct sites (K392 and K404) in the essential glycolytic enzyme enolase in Pseudomonas putida showcased the practical utility of our optimized GCE platform. In conclusion, we have successfully pioneered the establishment of acetylation in diverse bacterial strains, developed a host-agnostic GCE platform and identified strategies to optimize GCE efficiency in the hosting organism. These advancements herald new opportunities for studying PTMs in bacteria, unlocking numerous possibilities for both fundamental research and industrial biotechnology applications

## ACKNOWLEDGEMENTS

The research described in this paper is a part of the Predictive Phenomics Initiative at Pacific Northwest National Laboratory and conducted under the Laboratory Directed Research and Development Program. Additional support was provided by the U.S. Department of Energy (DOE), Office of Biological and Environmental Research (BER), as part of BER’s Genomic Science Program (GSP), and is a contribution of the Pacific Northwest National Laboratory (PNNL) Secure Biosystems Design Science Focus Area “Persistence Control of Engineered Functions in Complex Soil Microbiomes”. Pacific Northwest National Laboratory is a multiprogram national laboratory operated by Battelle for the U.S. Department of Energy under Contract No. DE-AC05-76RL01830. We also thank Adam Guss for sharing the SAGE-compatible *Pseudomonas putida* strain AG5577 and Alan Wolfe for insightful conversations regarding lysine acetylation in bacterial proteins.

## MATERIALS AND METHODS

### Reagents and media

All commercial chemicals are of reagent grade or higher. All solutions were prepared in deionized water that was further treated by Barnstead Nanopure^®^ ultrapure water purification system (Thermo Fisher Scientific). *Pseudomonas putida* AG5577, a derivative of KT2440 with multiple *attB* sites for compatibility with SAGE, was provided by Adam Guss. The strain was routinely cultured at 30 °C on Lysogeny Broth (LB) agar plates or in LB liquid medium. Antibiotics were added where appropriate to following final concentrations: kanamycin sulfate (50 μg/ml), apramycin sulfate (50 μg/ml), carbenicillin (50 μg/ml), gentamicin sulfate (15 μg/ml), neomycin sulfate (10 μg/ml), or spectinomycin/streptomycin (200/300 mg/L, respectively). Non-canonical amino acids 4-azido-L-phenylalanine (AzF), Acetyl-L-lysine (AcK) and N6-((Prop-2-yn-1-yloxy)carbonyl)-L-lysine hydrochloride (PlK) were purchased from Chem Impex, 4-Fluoro-L-phenylalanine (4FF), *p*-chlorophenylalanine (4ClF), 4-benzoyl-L-phenylalanine (BzF), *p*-Acetylphenylalanine and 4-Propargyloxy-L-phenylalanine were purchased from Cayman Chemicals and N6-((2-Azidoethoxy)carbonyl-L-lysine was purchased from Sigma Aldrich. Aromatic ncAAs were dissolved in equimolar NaOH to make 100 mM stock solutions. Lysine derivative ncAAs were dissolved in ddH2O to make 200 mM stock solutions. Solutions of antibiotics and ncAAs were filtered through 0.22 μm sterile membrane filters.

### General culturing conditions and media

The plasmids, strains and primers used in this study are listed in Tables S4-S6 respectively. Routine cultivation of *E. coli* for plasmid construction and maintenance was performed at 37°C using LB (Lennox) medium supplemented with antibiotics. LB (Lennox) was used for routine pseudomonad strain maintenance and competent cell preparations. LB (Lennox) or R2A medium was used for routine *R. jostii* strain maintenance and competent cell preparations. All solid medium was prepared by addition of agar (15 g/liter). MOPS (3-(N-morpholino)propanesulfonic acid))-based mineral medium (MME) supplemented with 10 mM NH4Cl and with 10 mM glucose were used for pseudomonad GCE experiments. MME contains 9.1 mM K_2_HPO_4_, 20 mM MOPS, 4.3 mM NaCl, 20 mM NH_4_Cl, 0.41 mM MgSO4, 68 μM CaCl_2_, and 1× MME trace minerals (pH adjusted to 7.0 with KOH). MME trace mineral stock solution (1000X) contains per liter 1 mL of concentrated HCl, 0.5 g of Na_4_EDTA, 2 g of FeCl_3_, and 0.05 g each of H_3_BO_3_, ZnCl_2_, CuCl_2_⋅2H_2_O, MnCl_2_⋅4H_2_O, (NH_4_)_2_MoO_4_, CoCl_2_⋅6H_2_O, and NiCl_2_⋅6H_2_O. An MME variant (MMV) additionally containing 10 mg each of niacin, pantothenate, lipoic acid, p-aminobenzoic acid, thiamine (B1), riboflavin (B2), pyridoxine (B6), and cobalamin (B12) and 4 mg each of biotin and folic acid per liter was used for *R. jostii* assays. All pseudomonad and *R. jostii* cultures were incubated at 30°C unless otherwise indicated. Cultures were incubated in 24-well deep well plates with shaking at 700 rpm in a heated plate shaker (Southwest Science).

### Plasmid construction

Q5 High-Fidelity DNA Polymerase (New England Biolabs (NEB)) and primers synthesized by Eurofins Genomics were used in all PCR amplifications for plasmid construction. OneTaq DNA polymerase (NEB) was used for colony PCR. EvaGreen dye (Biotium) was supplemented at 1× final concentration in PCRs for quantitative PCR (qPCR) applications. Plasmids constructed by Gibson Assembly using the NEBuilder HiFi DNA Assembly Master Mix (NEB) or ligation using T4 DNA ligase (NEB). Plasmids were transformed into competent NEB 5-alpha F’I^q^ (NEB). Standard chemically competent *E. coli* transformation protocols were used to construct plasmid host strains. Transformants were selected on LB (Lennox) agar plates containing antibiotics for selection and incubated at 37°C. Template DNA for PCRs was synthesized by Twist Biosciences. The Zymoclean Gel DNA recovery kit (Zymo Research) was used for all DNA gel purifications. Plasmid DNA was purified from *E. coli* using the GeneJet plasmid miniprep kit (Thermo Fisher Scientific) or the ZymoPURE II plasmid midiprep kit (Zymo Research). Sequences of plasmids were confirmed using Sanger sequencing performed by Eurofins Genomics or nanopore sequencing by Plasmidsaurus. Plasmids used in this work are listed in Table S4, and maps of plasmids constructed in this study can be found in Supplementary File P1.

### Strain construction

All genome modifications were performed by integration of non-replicating plasmids into the chromosome with serine recombinases via SAGE, where Bxb1, R4, or TG1 integrases unidirectionally recombine DNA between cognate attB and attP sequences. Plasmids containing GCE machinery harbor either a Bxb1, R4 or TG1 attP sequence, and variants of *Pseudomonas putida* KT2440 (AG5577), *Pseudomonas fluorescens* SBW25 (JE4621), *Pseudomonas fredericksbergensis* TBS10 (JE5041), *Pseudomonas facilor* TBS28 (RS175) and *Rhodococcus jostii* RHA1 have each been engineered to harbor the cognate Bxb1, R4 or TG1 attB sequence.

Plasmid transformations were performed using electroporation with electrocompetent cells. Electrocompetent cells were prepared by growing cultures as previously described^35^, in LB or R2A broth (for pseudomonads or RHA1 respectively) until cultures reached stationary phase. Fresh medium was inoculated with the preculture to an OD_600_ of 0.01 and incubated aerobically until the cells reached a mid-log phase (∼OD_600_ = 0.4 to 0.6). Once the desired culture density was reached, the cells were centrifuged at 5000*g* for 20 min at 4°C and washed at one-half culture volume of ice-cold 10% glycerol. Centrifugation and washing were repeated two additional times, and then cells were resuspended in 1/50th culture volume of ice-cold 10% glycerol. Competent cells were either used immediately or stored at −80°C.

Electroporation was performed as follows. GCE machinery and SAGE integrase plasmids (200 ng each) were added to 70 μL of ice-cold electrocompetent cells with the total volume of DNA not exceeding 5 uL. Competent cells were transferred into an ice-cold 0.1-cm-gap electroporation cuvette and electroporated at 1.6 kV, 25 μF, and 200 ohms. Following electroporation, 950 μL of SOC or R2A were added to the competent cells, and the resulting mixture was incubated aerobically, at 30°C with shaking for 3 hours. Following this recovery step, various dilutions of recovery cultures were plated on selective LB or R2A solid medium and incubated aerobically at 30°C. Primers used to screen for integration of the poly-*attP* cassette insertions in pseudomonads and *R. jostii* can be found in Table S6. Verified strains were stored at −80 C with 20% glycerol.

### GCE incorporation efficiency assays

Starter cultures for pseudomonads and RHA1 were prepared by inoculating 6 mL of LB or R2A media lacking antibiotic with appropriate strains from glycerol stocks. These strains were cultivated to stationary phase at 30°C with shaking until the cultures reached stationary phase. Cultures were then diluted 100-fold into 6 mL of MME with 10 mM NH4Cl and 10 mM glucose and incubated again at 30°C with shaking until the cultures gain reached saturation. The saturated cultures were used to inoculate GCE efficiency assays. All GCE assays were performed in 24-well deep well plates with shaking at 30°C (700 rpm, 1-mm orbital). The media used in assays (MME-GCE) was composed of MME with 10 mM NH4Cl, 10 mM glucose, and a 1× 18-amino acid solution^64^. Each strain was tested in quadruplicate under two conditions: with and without ncAA supplementation (+/− ncAA). For -ncAA cultures, no additional ncAA was added. For +ncAA cultures, either 1 mM ncAA for aromatic ncAAs or 10 mM ncAA for lysine derivatives were added to the media. Saturated cultures prepared as described above were used to inoculate 2mL of the assay media to obtain a 100-fold dilution. Cultures were incubated for 24 hours, after which cultures were diluted 100-fold in 1× PBS, and ncAA-sensitive fluorescence was measured using a NovoCyte flow cytometer (Acea Biosciences). Green fluorescence (sfGFP) was measured using a 488-nm laser with a 530/30-nm filtered detector. A preliminary gating of 2500 and 10 were used for forward scatter (FSC) and side scatter (SSC), respectively, and a total of 25,000 events were measured for each sample.

### Creation of Promoter-RBS library

A combination of 449 synthetic and natural promoter sequences (Table S1) were synthesized with the following parameters and coupled with 1 of 106 synthetic ribosomal binding sites designed by the RBS calculator (Table S2). Synthetic promoters were derived from commonly used promoters in the field of synthetic biology, as well as custom promoters generated for this study. Each promoter was coupled with a downstream synthetic RBS, designed by the RBS calculator to span a 441-fold range of translation initiation rates, during the cloning process. Promoter sequence DNA, as well as flanking sequences used for cloning, were synthesized as a pool of oligos by Twist Biosciences. To generate a *Mj*TyrRS expression variant, plasmid library primers oPNL2125 / oPNL2126 were used to amplify the backbone of pJE2027, excising the existing tac promoter and RBS. The promoter library was amplified using forward primer oPNL2127 and an equimolar amount of six reverse primers that each contain a priming sequence that binds to the oligo pool, one of six sets of degenerate RBS sequences, and a region of homology with the plasmid backbone. The plasmid backbone and promoter/RBS pool were amplified using Q5 High-Fidelity DNA Polymerase (NEB). PCR products were assembled using NEBuilder HiFi DNA Assembly Master Mix (NEB) and transformed into chemically competent NEB 5-alpha F’IQ competent cells and recovered in 1 mL of SOC at 37 °C for 1 hour. A 10-μl aliquot of the recovery mixture was diluted and plated to determine the cloning efficiency and library coverage, and the remaining 990 μL was propagated through overnight liquid selections using 50 ml LB + apramycin sulfate (50 μg/ml) grown at 37 °C, 200 rpm. The resulting plasmid library contained ∼209,000 transformants. Plasmid DNA was then extracted from library cultures with a ZymoPURE II Midiprep kit (Zymo Research) for subsequent transformation into final the host strains.

The plasmid pool was integrated into the R4 *attB* site in the chromosome of strain AG5577 containing a 2X tRNA cassette (pEVF108) by co-electroporation of 500 ng of the plasmid pool with 3 ug of pGW39 (R4 integrase expression plasmid) into 50 µL of electrocompetent cells. The electroporated cells were resuspended in 950 µL of SOC and incubated at 30°C for 1 hour with shaking. As above, dilutions generated from 10 μL of the recovery culture were plated onto LB plates supplemented with apramycin sulfate (50 μg/ml), and resulting colonies were quantified to identify the size of the library. The remaining 990 μL was propagated through overnight liquid selections using 50 ml LB + apramycin sulfate (50 μg/ml) grown at 30 °C, 200 rpm. Library cultures were diluted to OD_600_ = 1 with glycerol to a final concentration of 12.5% glycerol and stocked as 1-ml aliquots for use in subsequent experiments.

### Identification of active GCE-SAGE system with FACS-based selection

A FACS-based strategy was used to enrich and refine the *P. putida* promoter library based on sfGFP expression. The promoter library was introduced into *P. putida* KT2440 via electroporation as described above and glycerol stocks of the library were revived for selections. This was done by inoculating 50 mL of LB media with one 1 mL aliquot of the prepared glycerol stocks and incubating at 30°C for at least 6 hours. After, the culture was washed in MME (1X, 4 min at 5000 rcf) and resuspended in 50 mL MME with 10 mM glucose and incubated overnight at 30°C overnight.

This overnight culture was used to inoculate a 24-well deep well plate containing 2 mL MME-GCE with 10 mM NH4Cl, 10 mM glucose, and a 1× 18-amino acid solution as described above. This block contained wells with and without 1 mM pAzF for negative and positive selections of the library. The plate was also inoculated with strains harboring GCE v2 in the presence and absence of ncAA to provide fluorescent benchmarks for gating negative and positive events. Lastly the plate contained the parent strain AG5577 to establish background fluorescence. The plate was incubated overnight with shaking at 30°C (800 rpm, 1-mm orbital).

#### Positive selections

A Sony SH800 cell sorter was used for all sorting experiments with a four-laser set up (405, 488, 561, and 638 nm) and a 100-um sorting chip. The samples were sorted using the “Normal” sort mode into two-way 1.5 mL tubes using a forward scatter (FSC) threshold value of 0.05%. Initial gates were established on forward scatter—area by back scatter—area (FSC-A x BSC-A) and then forward scatter—height by forward scatter—width (FSC-H and FSC-W) to select for cells of expected size. The third gate was established with both the parent strains and the -ncAA condition for GCE v2.0 as a low-fluorescence control. Lastly, a fourth gate was drawn using the +ncAA condition to map out the range of GCE-produced fluorescence.

With these gates, the library incubated in the presence and absence of ncAA was analyzed. A ncAA-insensitive high fluorescence peak was observed in both populations, indicative of library members that may dysregulate GCE systems and charge native amino acids. A gate capturing the top 1.5% of events for the +ncAA library but avoiding the ncAA-insensitive events was drawn, and 10,000 of those events were bulk-sorted into a 1.5 mL Eppendorf tube for a positive selection. 1 mL of LB media was added to the tube and the positive library was incubated at 30°C to recover for two hours. After, 50 mL of MME with 10 mM glucose was inoculated with the recovered library and cultured for 48 hours at 30°C.

#### Negative selections

After incubation, the positive library was sub-cultured into fresh MME with 10 mM glucose at a 50-fold dilution and incubated overnight with shaking at 30°C. This culture was analyzed by the Sony SH800 cell sorter using gates drawn for low background events from the parent strain and GCE v2.0 in the -ncAA condition. Low background events were single cell sorted with high purity into a 96 well plate containing 150 uL of LB media. Controls, including the parent strain, and GCE v2.0 were also added to the plate to provide direct comparisons during validation. The plate was incubated with shaking at 30°C overnight, then used to inoculate a 96-deep well plate for glycerol stocks as well as a 96 well plate containing 150 uL of MME with 10 mM glucose for validation assays.

#### Validation experiments

After incubation overnight at 30°C with shaking, the above plate was used to inoculate two additional black-walled, clear-bottom 96-well microplates, one containing 150 uL of MME-GCE without ncAA to determine background incorporation of the potential hits and one with MME-GCE containing 1 mM pAzF to determine GCE efficiency. These plates were incubated for 24 hours using the “fast shaking” 1 mM double orbital setting on a Neo2SM plate reader (BioTek). Fluorescence (*F*_501,520_ for sfGFP) was then measured for both plates. Background fluorescence readings from wells containing the non-fluorescent parent strain were averaged and subtracted from sample readings before analysis. A subset of potential hits from the initial block were then screened in a typical GCE experiment using 2 mL of culture (see methods above) and promising clones were sequenced.

### Expression and purification of ncAA-protein for mass spectrometry analysis

To produce sufficient protein for mass spec validation, 5 mL LB cultures were inoculated from frozen glycerol stocks and cultured overnight at 30°C. Overnight cultures were subcultured into 5 mLs MME with 10 mM NH4Cl and 10 mM glucose and cultured overnight at 30°C. These cultures were used to inoculate 200 mL of MME-GCE with ncAA in a 1 L Erlenmeyer flask at a 1:100 dilution. These cultures were grown overnight at 30°C with shaking. The next day, the cultures were pelleted by centrifugation (15 min, 5500 rcf) and spent media was removed. The cell pellets were resuspended in a lysis/wash buffer (50 mM Tris-HCl, 500 mM NaCl, and 5 mM imidazole (pH 7.5) and lysed via sonication. Cell debris was pelleted at 15,000 rcf for 25 min at 4°C and clarified cell lysate was recovered. To bind His_6_-tagged protein, clarified cell lysate was incubated with 200 μL of Ni-NTA resin (HisPur™ Ni-NTA Resin, Thermo Scientific) at 4°C for 1 hour with rocking. Resin was collected and extensively washed with 50 resin bed volumes (bv) of lysis buffer. Bound protein was eluted from the resin by incubation with 5 bed volumes of elution buffer (lysis buffer with 200 mM imidazole) concentrated, and exchanged into storage buffer (50 mM phosphate, 150 mM NaCl) with a 3-kDa MWCO Vivaspin spin-concentration filter (GE Health Sciences). Protein concentration was determined by absorbance at 280 nm, flash-frozen with liquid nitrogen and stored at −80°C until needed. For purification of sfGFP_2XpAzF_ an additional size exclusion chromatography step was added (Superdex 200 increase 10_300 column from Cytiva Product #28990944, PBS Buffer pH 7.4, Flow rate 0.5mL/min). However, during analysis, sfGFP_2XpAzF_ experienced some proteolytic degradation, though maintained its fluorescent signature (Fig. S2).

For mass analysis of protein, the purified proteins were then analyzed by Creative Proteomics facility on a Synapt G2-Si mass spectrometer by the following method. The protein of interest was isolated using chromatographic techniques (C4 on the H-Class UPLC) and introduced to the Synapt G2-Si using a dual orthogonal API source (Lock-Spray ESI/APCI) and subjected to CID fragmentation. The mass spectrometer was operated in ESI positive resolution mode in the range 300-5000 Da. The capillary voltage was set to 3.0 kV with a sampling cone voltage of 78 V, a source temperature of 150°C and a desolvation temperature of 550°C. The cone gas was set to 0 L/h with desolvation gas at 300 L/h and the nebulizer set to 6.0 bar. The protein’s molecular weight can then be determined by deconvoluting the fragment masses using Waters MaxEnt software.

## Contributions

E.M.V.F and M.S. created the plasmids and strains with help from S.A and A.W. Assays were performed by M.S. with guidance from E.M.V.F and with help from S.A. Library analysis was performed by J.E. with help from A.W. Protein expression and purification was performed by M.S. with help from R.W. Protein modelling was done by Y.Y. The manuscript was mainly prepared by E.M.V.F. and with input from all authors. E.M.V.F, E.N, R.E. and J.E. supervised the research.

## ETHICS DECLARATIONS

### Competing interests

The authors declare no competing interests.

## Notes

### Competing Interest Statement

The authors have declared no competing interest.

### Summary of Updates

Text has been polished and figure legends updated.

