## Supplemental Materials for "Efficient genetic code expansion tools enable *in vivo* study of lysine acetylation in non-model bacteria"

**The file includes:**

Figs. S1 to S5

Tables S4 to S6

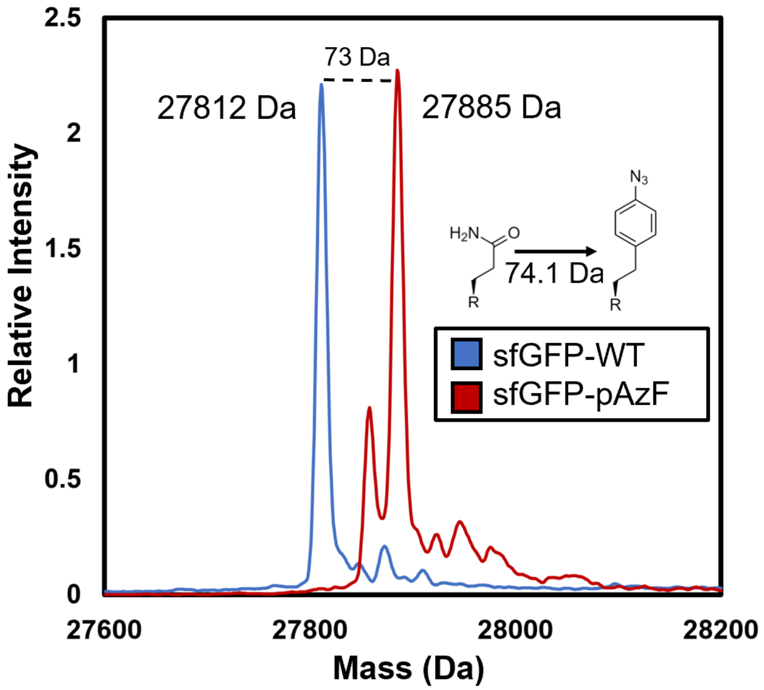


**Supplemental Fig. 1** Electrospray ionization mass spectrometry analyses of $sfGFP_{WT}$ (blue trace) and $sfGFP_{pAzF}$ (red trace). Observed masses are shown and correspond well to the expected masses (27811 Da and 27885 Da respectively) and the change in side chain at site 150 (N versus pAzF) is shown with the expected mass shift.

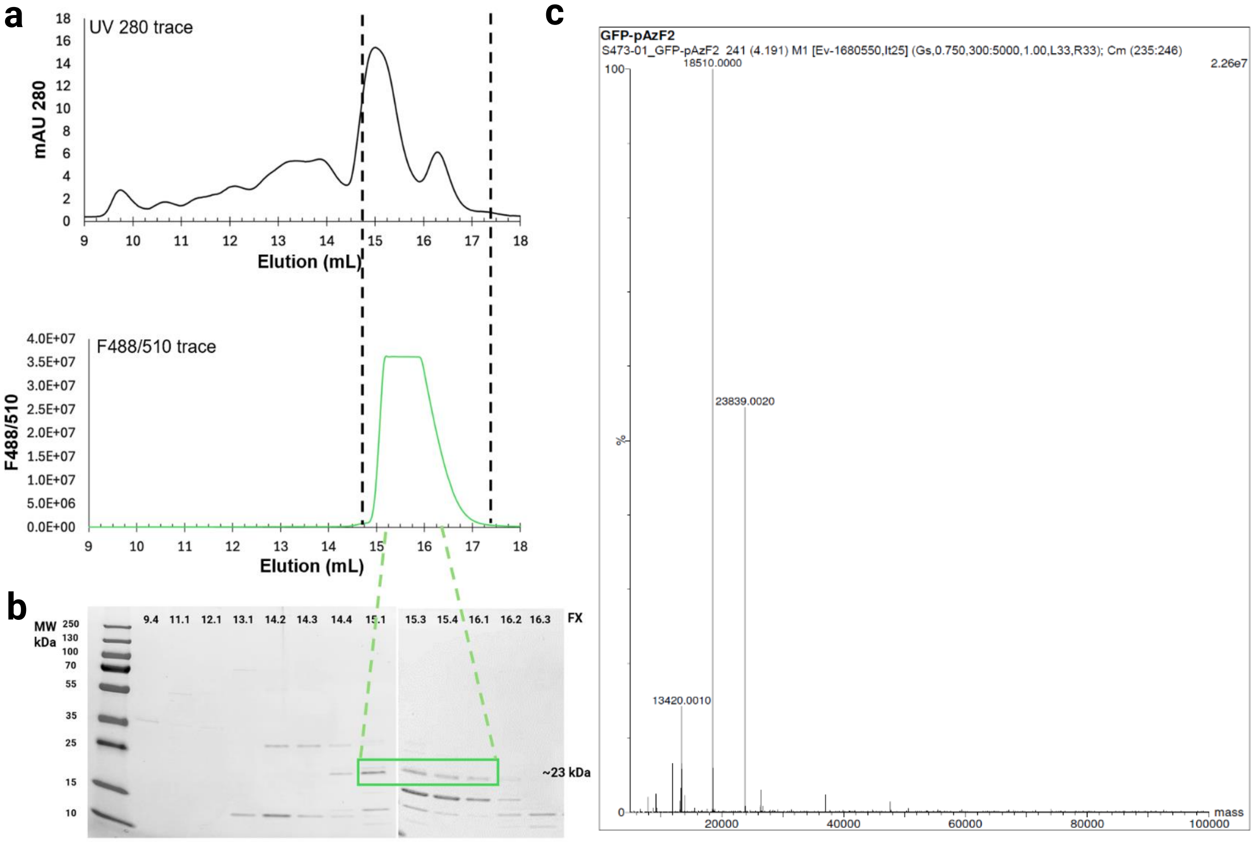


**d**

| Sequence of sfGFP | Calc. MW (Da) | Obsv. MW (Da) | Diff. (Da) |
|---|---|---|---|
| MRKDEELFTGVVPILVELDGDVNGHKFSVRGEGEGDATNGKLTLKFICTTGKLPVP WPTLVTTLTYGVQCFARYPDHMKQHDFFKSAMPEGYVQERTISFKDDGTYKTRAE VKFEGDTLVNRIELKGIDFKE**pAzF**GNILGHKLEYNFNSH**pAzF**VYITADKQKNGIKANF KIRHNVEDGSVQLADHYQQNTPIGDGPVLLPDNHYLSTQSVLSKDPNEKRDHMVLL EFVTAAGITHGMDELYKGSHHHHHH | 23840.9 | 23839 | 1.95 |

**Supplemental Fig. 2. a** UV 280 and F488/510 traces of size-exclusion chromatography of $sfGFP_{2XpAzF}$ (top and bottom panels respectively) **b** SDS-PAGE analysis of fractions from size-exclusion chromatography. Proteolytically degraded $sfGFP_{2XpAzF}$ with associated fluorescent signature is indicated in green box. **c** Electrospray ionization mass spectrometry analysis of proteolytically degraded $sfGFP_{2XpAzF.}$ **d** Sequence of $sfGFP_{2XpAzF,}$ likely cleaved residues are indicated in red. Calculated mass of degraded $sfGFP_{2XpAzF}$ corresponds well to the observed masses.

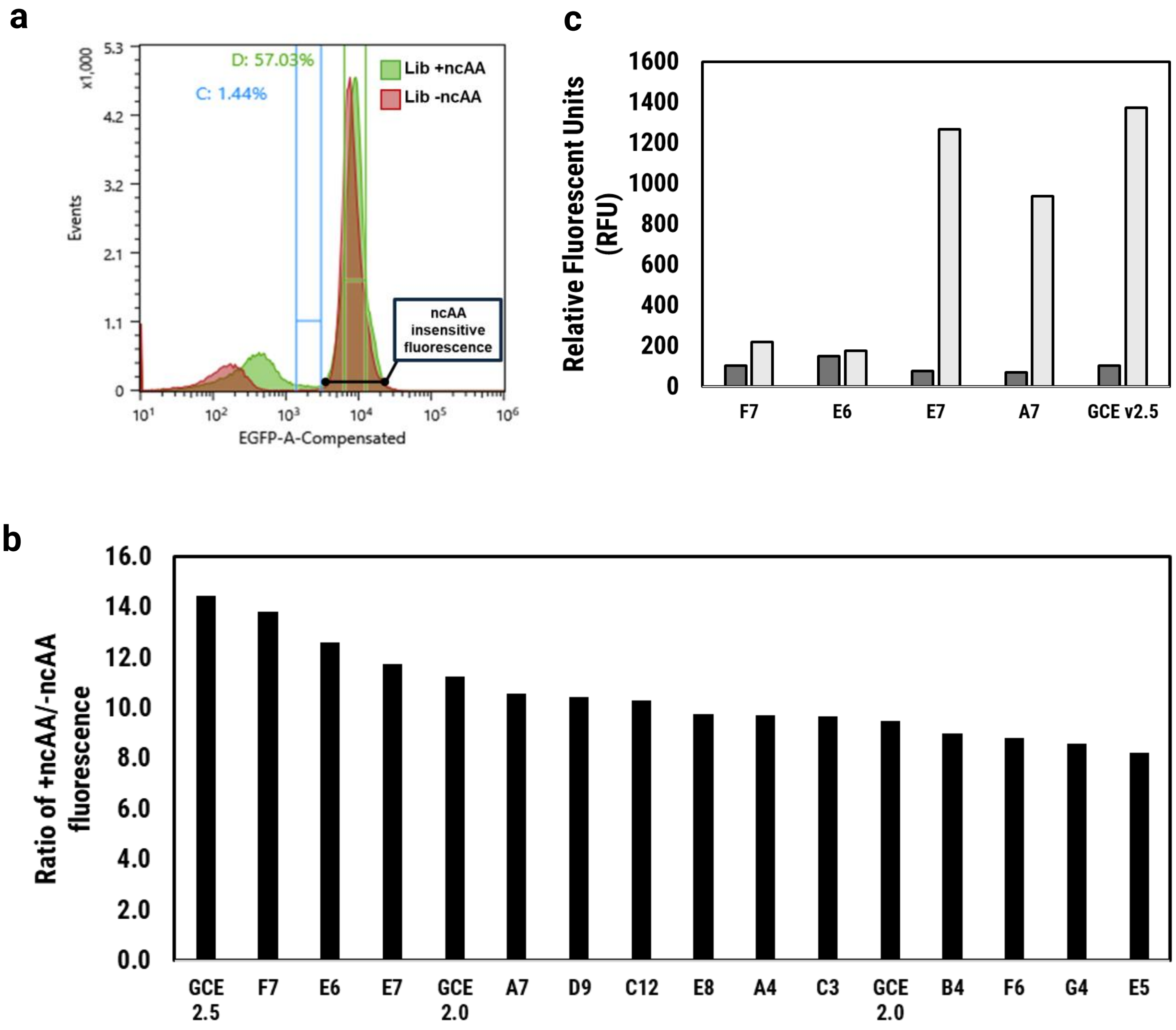


**Supplemental Fig. 3. a** Representative histograms of the flow cytometry analysis of the GCE-promoter library cultured with (green trace) and without (red trace) ncAA. The blue gate represents the population that was selected during positive sorts. The green gate represents a highly fluorescent ncAA insensitive population. **b** Results of 0.2 mL GCE incorporation efficiency assays on a subsection of top hits from 96 well single sort. The ratio of +ncAA/-ncAA is given on the y-axis. GCE v2.5 and GCE v2.0 were included on the plate to provide a direct comparison. **c** Results of 2 mL GCE incorporation efficiency assays used to identify final hits.

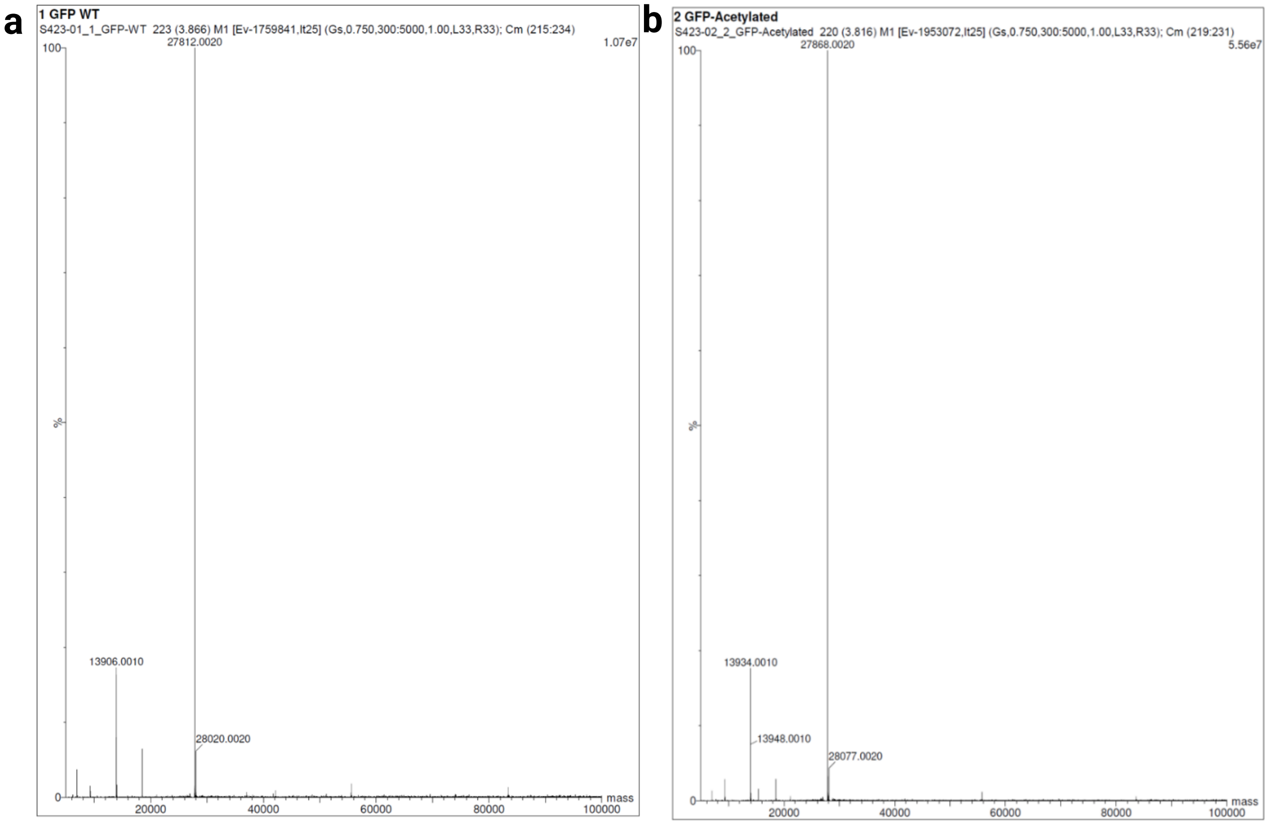


**Supplemental Fig. 4. a** Electrospray ionization mass spectrometry analyses of sfGFP. Observed masses are shown and correspond well to the expected mass (27811.3 Da) **b** Electrospray ionization mass spectrometry analyses of $sfGFP_{AcK150}$. Observed mass is shown and corresponds well to the expected mass (27867.4 Da)

a

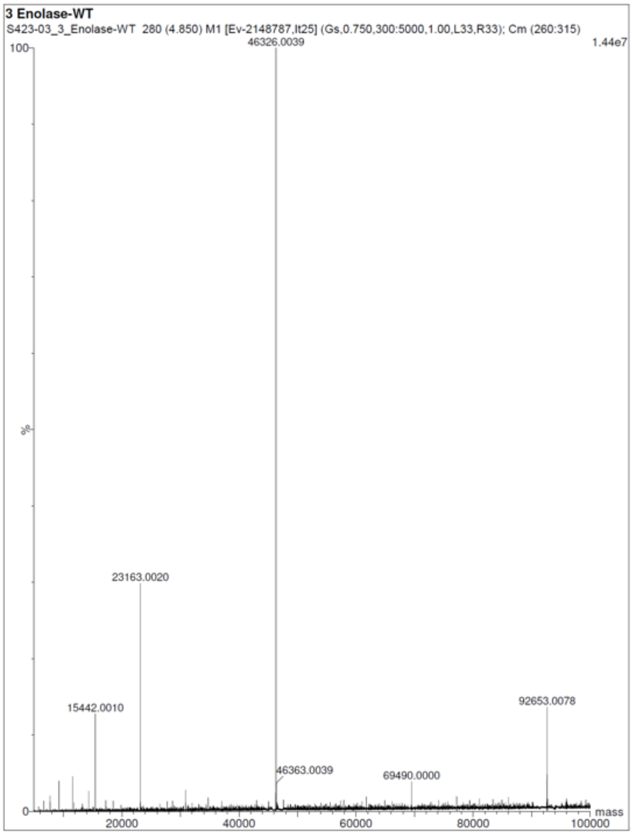


**Supplemental Fig. 5. a** Electrospray ionization mass spectrometry analyses of enolase$_{WT}$. Observed mass is shown. Expected mass is 46326.4 Da, including the loss of methionine. **b** Electrospray ionization mass spectrometry analyses of enolase$_{AcK342}$. Observed mass is shown. Expected mass is 46368.4 Da, including the loss of methionine. **c** Electrospray ionization mass spectrometry analyses of enolase$_{AcK404}$. Observed mass is shown. Expected mass is 46368.4 Da, including the loss of methionine.

b

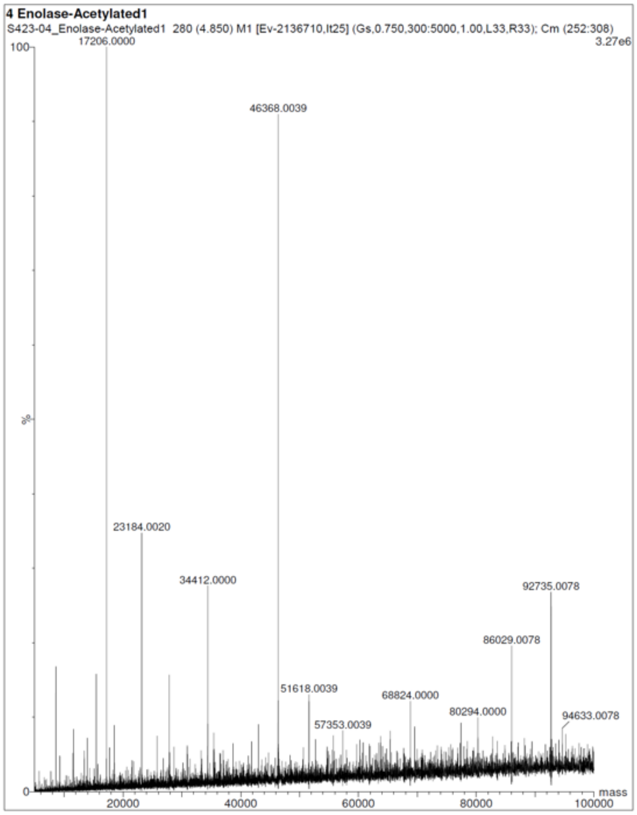


c

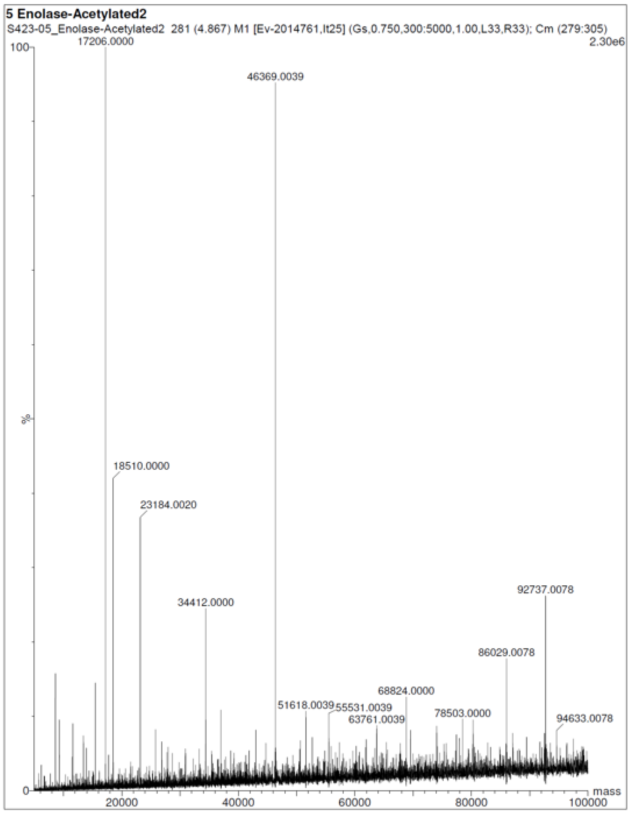

**Table S4 - Plasmids used in this study**

| Name | Genotype | Source |
|---|---|---|
| pJH225 | CloDF13, *aadA1* (spec/strep), R4 *attP* | This work |
| pJH219 | ColA, *aac(3)-I* (gent), TG1 *attP* | This work |
| pJH204 | ColE1, *nptII* (kan/neo), Bxb1 *attP* | This work |
| pAW45 | CloDF13, *aac(3)IV* (apr), R4 *attP* | This work |
| pEVF1 | pJH0225 JEa3:sfGFP_150 | This work |
| pEVF3 | pJH219 Leu-RS:Mj_TyrRS | This work |
| pEVF7 | pJH0204 Leu-tRNA:Mj_tRNA(1X) | This work |
| pEVF12 | pJH0225 JEa3:sfGFP_WT | This work |
| pEVF45 | pAW45 JEa3:sfGFP_WT | This work |
| pEVF46 | pAW45 Leu-RS:Mj_TyrRS, JEa3:sfGFP_150 | This work |
| pEVF75 | pJH0204 Leu-tRNA:Mj_tRNA(2X),EF-Tu | This work |
| pEVF108 | pJH0204 Leu-tRNA:Mj_tRNA(2X) | This work |
| pEVF148 | pAW45 Leu-RS:Mj_TyrRS, JEa3:sfGFP_134,150 | This work |
| pMS15 | pJH0204, nptll, Ala-tRNA:Mj_tRNA(2X),EF-Tu | This work |
| pMS3 | pAW45 Ala-RS:Mj_TyrRS_CO, Cym:sfGFP_150_CO | This work |
| pMS16 | pAW45 Ala-RS:Mj_TyrRS, Cym:sfGFP_150 | This work |
| pJE2027 | pAW45 Ptac:Mj_TyrRS, JEa3:sfGFP_150 | This work |
| pEVF210 | pAW45 A7:Mj_TyrRS, JEa3:sfGFP_150 | This work |
| pEVF211 | pAW45 E7:Mj_TyrRS, JEa3:sfGFP_150 | This work |
| pEVF212 | pAW45 F7:Mj_TyrRS, JEa3:sfGFP_150 | This work |
| pEVF109 | pJH0204 Leu-tRNA:Pyl_tRNA(2X) | This work |
| pEVF145 | pAW45 Leu-RS:chimAcK3(IPYE), JEa3:sfGFP_150 | This work |
| pEVF146 | pAW45 Leu-RS:MbAcK3(IPYE), JEa3:sfGFP_150 | This work |
| pEVF190 | pAW45 Leu-RS:chimAcK3(IPYE), JEa3:enolase_WT | This work |
| pEVF191 | pAW45 Leu-RS:chimAcK3(IPYE), JEa3:enolase_TAG392 | This work |
| pEVF192 | pAW45 Leu-RS:chimAcK3(IPYE), JEa3:enolase_TAG404 | This work |

**Table S5 - Strains used in this study**

| Name | Genotype | Source |
|---|---|---|
| NEB 5-alpha F'Iq | Escherichia coli F´ proA+B+ lacIq Δ(lacZ)M15 zzf::Tn10 (TetR) / fhuA2Δ(argF-lacZ)U169 phoA glnV44 Φ80Δ(lacZ)M15 gyrA96 recA1 relA1 endA1 thi-1 hsdR17 | New England Biolab |
| KT2440 | *Pseudomonas putida* KT2440 | *(35)* |
| SBW25 | *Pseudomonas fluorescens* SBW25 | *(44)* |
| TBS28 | *Pseudomonas facilor* TBS28 | *(45)* |
| TBS10 | *Pseudomonas frederiksbergensis* TBS10 | *(31)* |
| RHA1 | *Rhodococcus jostii* RHA1 | *(46)* |
| AG5577 | *P. putida* KT2440::10x poly-attB | *(31)* |
| JE4621 | *P. fluorescens* SBW25 ampC:10x poly-attB | *(31)* |
| RS175 | *P. facilor*::10x poly-attB | *(45)* |
| JE5041 | *P. frederiksbergensis*::10x poly-attB | *(31)* |
| AG5879 | *R. jostii* RHA1_RS20555::10x poly-attB | *(31)* |
| EVF7 | *P. putida* AG5577 *attL*R4:pEVF1: *attR*R4, *attL*Bxb1:pEVF7*: attR*Bxb1, attLTG1:pEVF3: attRTG1 | This work |
| EVF12 | *P. putida* AG5577 *attL*R4:pEVF45: *attR*R4 | This work |
| MS5 | *P. putida* AG5577 *attL*R4:pEVF105: *attR*R4, *attL*Bxb1:pEVF108*: attR*Bxb1 | This work |
| EVF44 | *P. putida* AG5577 *attL*R4:pEVF105: *attR*R4, *attL*Bxb1:pEVF75*: attR*Bxb1 | This work |
| MS44 | *P. putida* AG5577 *attL*R4*:pEVF105: attR*R4, *attL*Bxb1*:pEVF108: attR*Bxb1, attLTG1:pEVF141: attRTG1 | This work |
| MS36 | *R. jostii* AG5879 attLR4:pMS3: attRR4, attLTG1:pMS2: attRTG1 | This work |
| MS37 | *R. jostii* AG5879 attLR4:pMS16: attRR4, attLTG1:pMS2: attRTG1 | This work |
| MS61 | *P. fluorescens* JE4621 attLR4:pEVF105: attRR4, attLBxb1:pEVF75: attRBxb1 | This work |
| MS62 | P. frederiksbergensis JE5041 attLR4:pEVF105: attRR4, attLBxb1:pEVF75: attRBxb1 | This work |
| MS63 | *P. facilor* RS175 attLR4:pEVF105: attRR4, attLBxb1:pEVF75: attRBxb1 | This work |
| MS52 | *P. putida* AG5577 *attL*R4:pEVF145: *attR*R4, *attL*Bxb1:pEVF109*: attR*Bxb1 | This work |
| MS53 | *P. putida* AG5577 *attL*R4:pEVF145: *attR*R4, *attL*Bxb1:pEVF109*: attR*Bxb1, *attL*TG1:pEVF142: *attR*TG1 | This work |
| MS54 | *P. putida* AG5577 *attL*R4:pEVF146: *attR*R4, *attL*Bxb1:pEVF109*: attR*Bxb1 | This work |
| MS55 | *P. putida* AG5577 *attL*R4:pEVF146: *attR*R4, *attL*Bxb1:pEVF109*: attR*Bxb1, *attL*TG1:pEVF142: *attR*TG1 | This work |
| MS78 | *P. putida* AG5577 *attL*R4:pEVF190: *attR*R4, *attL*Bxb1:pEVF109*: attR*Bxb1, *attL*TG1:pEVF142: *attR*TG1 | This work |
| MS79 | *P. putida* AG5577 *attL*R4:pEVF191: *attR*R4, *attL*Bxb1:pEVF109*: attR*Bxb1, *attL*TG1:pEVF142: *attR*TG1 | This work |
| MS80 | *P. putida* AG5577 *attL*R4:pEVF192: *attR*R4, *attL*Bxb1:pEVF109*: attR*Bxb1, *attL*TG1:pEVF142: *attR*TG1 | This work |
| MS78 | *P. putida* AG5577 *attL*R4:pEVF190: *attR*R4, *attL*Bxb1:pEVF109*: attR*Bxb1, *attL*TG1:pEVF142: *attR*TG1 | This work |
| MS79 | *P. putida* AG5577 *attL*R4:pEVF191: *attR*R4, *attL*Bxb1:pEVF109*: attR*Bxb1, *attL*TG1:pEVF142: *attR*TG1 | This work |
| MS80 | *P. putida* AG5577 *attL*R4:pEVF192: *attR*R4, *attL*Bxb1:pEVF109*: attR*Bxb1, *attL*TG1:pEVF142: *attR*TG1 | This work |

### Table S6 – Primers used in this study

| **Name** | **Sequence** |
|---|---|
| EVF_GCE_GFP_scrn_F | atgcgtaaagacgaagagctg |
| EVF_GCE_GFP_scrn_R | ctttgtacagttcatccataccatg |
| EVF_GCE_aaRS_scrn_F | ATGGATGAGTTTGAGATGATTAAACGC |
| EVF_GCE_aaRS_scrn_R | CAGGCGTTTGCGAATAGG |
| EVF_GCE_tRNA_scrn_F | GCTGCAGTGCATAAACAGC |
| EVF_GCE_tRNA_scrn_R | gaAAAGCTTTACATCATCTGCAGAAG |
| oPNL2125 | CGGATTGCAATTGAAGACTTGG |
| oPNL2126 | ATGGATGAGTTTGAGATGATTAAAC |
| oPNL2127 | CCAAGTCTTCAATTGCAATCCG |
| GCE_lib_v1_aaRS_RBSv1 | GTTTAATCATCTCAAACTCATCCATTTATTGHSCCCSYTTGCATTATTCGTTAAACAAAATTATTTGTAGAGGCTGT |
| GCE_lib_v1_aaRS_RBSv2 | GTTTAATCATCTCAAACTCATCCATTATTCGACSTCCTHTACCDCACACCTTAAACAAAATTATTTGTAGAGGCTGT |
| GCE_lib_v1_aaRS_RBSv3 | GTTTAATCATCTCAAACTCATCCATTTATTTRSCTCSTTTGCATTATTSCTTAAACAAAATTATTTGTAGAGGCTGT |
| GCE_lib_v1_aaRS_RBSv4 | GTTTAATCATCTCAAACTCATCCATTAGCAKACHKCCTTAACTGCAGCCSTTAAACAAAATTATTTGTAGAGGCTGT |
| GCE_lib_v1_aaRS_RBSv5 | GTTTAATCATCTCAAACTCATCCATTGTAAAACCKBCTTAACTGHAGCTTTTAAACAAAATTATTTGTAGAGGCTGT |
| GCE_lib_v1_aaRS_RBSv6 | GTTTAATCATCTCAAACTCATCCATAGATDSCCATCCCTAGTKCCGTGGGTTAAACAAAATTATTTGTAGAGGCTGT |
